# Cortical responses to natural speech reflect probabilistic phonotactics

**DOI:** 10.1101/359828

**Authors:** Giovanni M. Di Liberto, Daniel Wong, Gerda Ana Melnik, Alain de Cheveigné

## Abstract

Humans comprehend speech despite the various challenges of real-world environments, such as loud noise and mispronunciation. Our auditory system is robust to these thanks to the integration of the upcoming sensory input with prior knowledge and expectations built on language-specific regularities. One such regularity regards the permissible phoneme sequences, which determine the likelihood that a word belongs to a given language (phonotactic probability; “blick” is more likely to be an English word than “bnick”). Previous research suggested that violations of these rules modulate brain evoked responses such as the N400 and the late positive complex. Yet several fundamental questions remain unresolved, especially regarding the neural encoding and integration strategy of phonotactic information. Here, we used linear modelling approaches to assess the influence of phonotactic probabilities on the brain responses to narrative speech measured with non-invasive EEG. We found that the relationship between continuous speech and EEG responses is best described when the speech descriptor includes phonotactic probabilities. This provides us with a methodology to isolate and measure the brain responses to phonotactics using natural speech at the individual subject-level. Furthermore, such low-frequency signals showed the strongest speech-EEG interactions at latencies of 100-400 ms, supporting a pre-lexical role of phonotactic information.

**Significance Statement:** Speech is composed of basic units, called phonemes, whose combinations comply with language-specific regularities determining whether a sequence “sounds” as a plausible word. Our ability to detect irregular combinations requires matching incoming sequences with our internal expectations, a process that supports speech segmentation and learning. However, the neural mechanisms underlying this phenomenon have not yet been established. Here, we examine this in the human brain using narrative speech. We identified a brain signal reflecting the likelihood that a word belongs to the language, which may offer new opportunities to investigate speech perception, learning, development, and impairment. Our data also suggest a pre-lexical role of this phenomenon, thus supporting and extending current mechanistic perspectives.

## Introduction

Speech can be described as a succession of categorical units called *phonemes* that comply with language-specific regularities determining admissible combinations within a word. A sequence is said *well formed* if it sounds plausible as a word to native speakers (e.g. *blick*) and *ill formed* if it is perceived as extraneous to the language (e.g. *bnick*) (Chomsky and Halle, 1968; Parker, 2012). This concept is referred to as *phonotactics*. Well-formedness is gradient (Scholes, 1966; Chomsky and Halle, 1968; Frisch et al., 2000; Bailey and Hahn, 2001a; Hammond, 2004), meaning that we can assign a numerical value to each sequence of phonemes describing its likelihood of belonging to the language. Phonotactics aids lexical access (Vitevitch et al., 1999) and speech segmentation (Brent and Cartwright, 1996; Mattys et al., 1999) by constraining the space of likely upcoming phonemes, thus contributing to the robustness of speech perception to challenges such as noise, competing speakers, and mispronunciation (Davidson, 2006a; Obrig et al., 2016). High phonotactic probability facilitates learning of new words (Storkel and Rogers, 2000; Storkel, 2001, 2004; Storkel and Morrisette, 2002) and low phonotactic probability (violation) may trigger an attempt to repair a sequence into a well-formed word (Dehaene-Lambertz et al., 2000; Hallé et al., 2008; Carlson et al., 2016). However, considerable uncertainty remains about the cortical mechanisms underpinning the contribution of phonotactic information to speech comprehension (Winther Balling and Harald Baayen, 2008; Balling and Baayen, 2012; Ettinger et al., 2014). While part of the debate regards the pre- or post-lexical role of phonotactics, there is currently a lack of neurobiological data examining the cortical representation of phonotactic statistics. Hypotheses range from the explicit encoding of phoneme-level probabilities to the use of the lexical neighbourhood size as a proxy measure (McClelland and Elman, 1986; Bailey and Hahn, 2001b; Pisoni and Remez, 2005; Leonard et al., 2015).

One way to illuminate these issues is through the direct measurement of brain activity using technologies with high-temporal resolution, such as electroencephalography (EEG). Brain responses to phonotactics emerge by contrasting EEG responses to well- and ill-formed speech tokens, i.e. phonotactic mismatch response (PMM; Connolly and Phillips, 1994; Dehaene-Lambertz et al., 2000). This paradigm has been largely exploited in the literature, with somewhat sparse and inconsistent results. EEG responses to these violations emerge at latencies consistent with other well-known brain components, such as the mismatch-negativity (MMN), N400, and late positive complex (LPC) (Dehaene-Lambertz et al., 2000; Wiese et al., 2017). However, various types of confounds hamper the identification of responses specific to phonotactics. One issue is that brain responses to phonotactic probability may overlap with those reflecting subsequent processes, such as *learning* in case of novel well-formed sequences (pseudowords) and *phonological repair* for ill-formed tokens (non-words) (Bailey and Hahn, 2001a; White and Chiu, 2017). Secondly, if meaningful words are contrasted with ill-formed tokens, lexical-level N400 responses may arise that confound the contrast (Kutas and Federmeier, 2011; Rossi et al., 2011). The use of nonsense words avoids this issue, but the paradigm becomes more artificial. Natural speech may allow to investigate the cortical processing of phonotactics without such confounds; however it is generally characterised by well-formed words, therefore measuring PMM responses may be either not possible or suboptimal.

A novel approach to investigate the brain responses to natural speech may provide a solution to these issues (Di Liberto et al., 2015, 2018a; Crosse et al., 2016b; Broderick et al., 2018; de Cheveigné et al., 2018b). This method, based on linear modelling, allows to isolate and measure cortical responses to linguistic features of interest (e.g. phonemes) using natural speech stimuli. Here, we combine this approach with a computational model of phonotactics to test whether narrative speech elicits robust brain responses time-locked to patterns of phonotactic probabilities. We characterise the dynamics of cortical signals that are representative of real-life speech perception, contributing to the debate on the underpinnings of the cortical processes specific to phonotactics.

## Material and methods

The present study is based on new analyses of a previously published EEG dataset on natural speech perception (Di Liberto et al., 2015). The data include both the audio stimulus and the EEG response of the subjects listening to that stimulus. Data analysis involves fitting the EEG to various representations of the stimulus using a linear model. The quality of fit is used as an indicator of the relevance of each representation as a predictor of the cortical activity evoked in the listener by the speech stimulus.

### Subjects and Experimental Procedure

Ten healthy subjects (7 male) aged between 23 and 38 years old participated in the experiment. Participants reported no history of hearing impairment or neurological disorder. The experiment was carried out in a single session for each subject. Electroencephalographic (EEG) data were recorded from participants as they undertook 28 trials, each of ~155 seconds in length, where they were presented with an audiobook version of a classic work of fiction read by a male American English speaker. The trials preserved the storyline, with neither repetitions nor discontinuities. All stimuli were presented monophonically at a sampling rate of 44,100 Hz using Sennheiser HD650 headphones and Presentation software from Neurobehavioral Systems (http://www.neurobs.com). Testing was carried out in a dark room and subjects were instructed to maintain visual fixation for the duration of each trial on a crosshair centered on the screen, and to minimize eye blinking and all other motor activities. All procedures were undertaken in accordance with the Declaration of Helsinki and were approved by the Ethics Committees of the School of Psychology at Trinity College Dublin, and the Health Sciences Faculty at Trinity College Dublin. Further details about the stimulus and recording are available in Di Liberto et al., (2015) and the data is available at https://datadryad.org/resource/doi:10.5061/dryad.070jc.

### Speech representations

The approach used here follows a system identification framework that aims at disentangling brain responses to different speech and language features (Di Liberto et al., 2015). To this end, we first need to define such features (note that the first two elements are as in Di Liberto et al., 2015):

1. Acoustic spectrogram (**S**): This was obtained by filtering the speech stimulus into 16 frequency-bands between 250 Hz and 8 kHz distributed according to Greenwood’s equation (equal distance on the basilar membrane; Greenwood, 1961) using Chebyshev type 2 filters (order 100), and then computing the Hilbert amplitude envelope (the absolute value of the analytical signal obtained by the Hilbert Transform) for each frequency band.
2. Phonetic features (**F**): This multivariate representation of speech encodes phoneme-level information using phonetic features. The Prosodylab-Aligner software (Gorman et al., 2011) was used to partition each word into phonemes from the American English International Phonetic Alphabet (IPA) and align the speech stimulus with its textual transcription. This procedure returns estimates of the starting and ending time-points for each phoneme. Indicator functions for each of the 35 phonemes were recoded as a multivariate time series of 19 indicator variables, one for each of 19 phonetic features (based on the University of Iowa’s phonetics project http://soundsofspeech.uiowa.edu/) coding the manner of articulation (plosive, fricative, affricate, nasal, liquid, and glide), place of articulation (bilabial, labio-dental, lingua-dental, lingua-alveolar, lingua-palatal, lingua-velar, and glottal), voicing of a consonant (voiced and voiceless), and backness of a vowel (front, central, and back). Also, a specific feature was reserved for diphthongs. Each indicator variable took the value 1 between the start and the end of the phoneme (if relevant) and 0 elsewhere. Each phoneme was characterised by a value of 1 for some combination of indicator variables; not all such combinations map to permissible phonemes.
3. Phoneme onsets (**O**): This vector marks phoneme onsets with a discrete-time unit impulse, corresponding to the half-wave rectified first derivative of F. This is a non-linear transformation of the F features, thus linear models may benefit from the explicit definition of O combined with F.
4. Finally, we propose a novel representation using *phonotactic probabilities* (**P**). Natural languages include various constraints on the permissible phoneme sequences. Probabilities can be derived for a given speech token from this set of constraints. For example, the pseudoword *blick* would “sound” better than *bnick* to a native English speaker, which is reflected by a higher phonotactic probability for the first word. Here, we used a computational model (BLICK; Hayes and Wilson, 2008) based on a combination of explicit theoretical rules from traditional phonology and a maxent grammar (Goldwater and Johnson, 2003), which find the optimal weights for such theoretical constraints to best match the phonotactic intuition of a native speaker. Specifically, given a phoneme sequence ph_1..n_, P is composed of two vectors: a) inverse phonotactic probability (score(ph_1..n_) is the output of the BLICK software; it is small for well-formed tokens and large for ill-formed ones) and b) within-word derivative of the phonotactic probability (score(ph_1..(n-1)_) - score(ph_1..n_)), which describes the contribution of the latest phoneme to the well-formedness of the sequence.

In order to assess and quantify the contribution of each of the features F, O, and P to the speech-EEG mapping, the main analyses were conducted on the cumulative combinations S, FS, OFS, and POFS. The rationale is that, if the new feature carries information not subsumed by the other features, including it will improve the fitting score. To control for any potential effect of the difference in dimensionality of the feature space, we also used variants where the newly introduced feature did not correspond to the auditory stimulus, i.e. was shuffled (the entire procedure, including model fit, was rerun for each shuffled version). These mismatched vectors/matrices were generated by randomly shuffling: a) Phonetic features in the FS speech representation (F_shu_S) (every given phoneme, corresponding to a combination of N_F_ phonetic features, 1 for vowels and 3 for consonants, was replaced by N_F_ random phonetic features for its entire duration); b) Onset time in OFS (O_shu_FS) (the onset vector O was replaced by vector with the same number of impulses at random time points); and c) Phonotactic probability values in POFS (P_shu_OFS) (the values in the phonotactic vector P were randomly permuted while keeping the time information).

In addition to the phonotactic vector P, we defined three other representations that could reflect the encoding of phonotactic information in the brain. First, P_neigh_ is a vector of phoneme onsets amplitude-modulated using *neighborhood density* values. This information indicates the number of phonological neighbours given a speech token, where a phonological “neighbour” is a sequence of phonemes that can be obtained from the given token by deletion, addition, or substitution of a single phoneme. Similarly, P_sur_ and P_ent_ are vectors of phoneme onsets that are amplitude-modulated using phoneme *surprisal* and *entropy* respectively. These were calculated using the purely probabilistic measures “phoneme surprisal” and “cohort entropy” as defined by Gaston and Marantz (2018).

#### Phonotactic Probability Model

Phonotactic probability vectors were derived using the BLICK algorithm (Hayes and Wilson, 2008), a state-of-the-art tool based on explicit theories of phonology. Specifically, the BLICK algorithm constructs maxent grammars (e.g. Goldwater and Johnson, 2003) consisting of a set of numerically weighted phonological constraints. A training stage identifies weights that optimally match the phonotactic well-formedness intuition of experts. These weights are determined according to the principle of maximum entropy and, in the present work, were pre-assigned using an English grammar model (Hayes, 2012). Combining this pre-trained grammar with the textual transcription of the audio-book stimulus, BLICK performs a weighted sum of its constraint violations to calculate probability values reflecting the well-formedness of each speech token. Given a word, two scores were calculated for each phoneme token. The first indicates the inverse probability of the word segment up to that phoneme (e.g. the scores for /b/, /b l/, /b l ɪ /, and /b l ɪ k/ were calculated in correspondence of the four phonemes of the word ‘blick’). This time series of inverse probabilities was coded by the amplitudes of a series of pulses synchronous with those of the onset vector. The second is the finite difference of consecutive inverse probability values within a word (starting from the second phoneme of each word, e.g. P(/b/)–P(/b l/), P(/b l/)–P(/b l ɪ /), P(/b l ɪ /)–P(/b l ɪ k/); the score for the first phoneme of a word was assigned to the same value as in the phonotactic probability vector). The time series of difference measures was also coded as a time series of pulses synchronous with O. The concatenation of these two pulse trains constitutes the 2-dimensional phonotactic probability vector P.

### Data Acquisition and Preprocessing

Electroencephalographic (EEG) data were recorded from 128 scalp electrodes (plus 2 mastoid channels), filtered over the range 0 - 134 Hz, and digitised with a sampling frequency of 512 Hz using a BioSemi Active Two system. Data were analysed offline using MATLAB software (The Mathworks Inc.). EEG data were digitally filtered between 0.5 and 32 Hz using a Butterworth zero-phase filter (low- and high-pass filters both with order 2; implemented with the function *filtfilt*), and down-sampled to 64 Hz. EEG channels with a variance exceeding three times that of the surrounding channels were replaced by an estimate calculated using spherical spline interpolation (EEGLAB; Delorme and Makeig, 2004). All channels were then re-referenced to the average of the two mastoid channels with the goal of maximizing the EEG responses to the auditory stimuli (Luck, 2005).

#### Dimensionality reduction

The analyses that follow involve fitting the stimulus representation to the EEG response using a linear model. Both the stimulus and the EEG include a large number of dimensions (channels) many of which are correlated. To limit the risk of overfitting, it is useful to reduce their dimensionality. This is typically performed using principal component analysis (PCA). PCA finds a matrix of size N × N (if the data have N channels) that transforms the data to N ‘principal components’ (PC). The variance of the PCs sum up to the variance of the data. Subject to that constraint, the first principal component is the linear transform of the data with the largest possible variance. The second has the largest variance of transforms orthogonal to the first and so on. The first few PCs pack most of the variance, and so little variance is lost if a subset of N_PC_ < N PCs are selected and the remainder discarded. This procedure is applied repeatedly in the following analyses. In each case N_PC_ is tuned as a hyperparameter in a crossvalidation procedure to optimise the tradeoff between information retained and overfitting.

#### Denoising with multiway CCA

Our goal of evaluating the relevance of high-level speech structure representations by measuring their ability to predict cortical responses is hampered by the high level of noise and artifact in the EEG. We use a novel tool, multiway canonical correlation analysis (MCCA) to merge EEG data across subjects so as to factor out the noise. MCCA is an extension of canonical correlation analysis (CCA; Hotelling, 1936; de Cheveigné et al., 2018a) to the case of multiple (> 2) datasets. Given *N* multichannel datasets Xi with size T × Ji, 1 ≤ *i* ≤ *N* (time × channels), MCCA finds a linear transform Wi (sizes Ji × J0, where J0 < min(Ji)1 ≤ i ≤ N) that, when applied to the corresponding data matrices, aligns them to common coordinates and reveals shared patterns (de Cheveigné et al., 2018a). These patterns can be derived by summing the transformed data matrices: 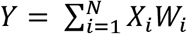. Thecolumns of the matrix Y, which are mutually orthogonal, are referred to as summary components (SC) (de Cheveigné et al., 2018a). Intuitively, the first few components are signals that most strongly reflect the shared information across the several input datasets. Here, these datasets are EEG responses to a same speech stimulus for 10 subjects.

This technique allows to extract a *consensus* signal that is shared across participants. The present study utilises this approach to test whether EEG responses to speech reflect phonotactic information. This methodology overcomes limitations of previous studies that attempted to obtain similar consensus responses by averaging data across subjects, which could not perform corregistration because of the lack of anatomical information and, therefore, ignored the likely topographical discrepancies between participants EEG signals (O’Sullivan et al., 2014; Di Liberto and Lalor, 2017). MCCA accounts for such discrepancies without the need for corregistration. Under the assumption that brain responses to speech share some fundamental similarities within a homogeneous group of normal hearing young adults, the MCCA procedure allows us to extract such common responses to the stimulus from other, more variable aspects of the EEG signals, such as subject-specific noise. For this reason, our analysis focuses on the first N_SC_ summary components, which we can consider as reflecting a ground truth EEG response to speech. N_SC_ was arbitrarily set to the number of dimensions for a single subject after dimensionality reduction (N_PC_; see the following section). This conservative choice was made by taking into consideration that the irrelevant signals within the retained components are excluded through the more restrictive CCA analysis that follows.

### Analysis Procedure

#### Stimulus-response model based on Canonical Correlation Analysis

Speech elicits brain responses that can be recorded with EEG. However, a large part of the EEG signal is unrelated to the stimulus as it may reflect other brain processes, as well as various forms of noise (e.g. muscle movements). Similarly, certain features of the speech input may have little or no impact on the measured brain responses. Studying the relation between speech and the corresponding EEG responses would greatly benefit from the ability to remove those unrelated portions of speech and EEG. This can be done by using canonical correlation analysis (CCA), a powerful technique that linearly transforms both stimulus and brain measurements so as to minimise irrelevant variance (Hotelling, 1936; de Cheveigné et al., 2018b).

In its more general definition, given two sets of multichannel data X_1_ and X_2_ of size T × J_1_ and T × J_2_, CCA finds linear transformations of both that make them maximally correlated. Specifically, CCA produces the transformation matrices W_1_ and W_2_ (sizes J_1_× J_0_ and J_2_ × J_0_, where J_0_ < min(J_1_,J_2_)) that maximise the correlation between pairs of columns of X_1_W_1_ and X_2_W_2_, while making the columns of each transformed data matrix X_i_W_i_ mutually uncorrelated. The first pair of canonical components (CC) is the linear combination of X_1_ and X_2_ with highest possible correlation. The next pair of CCs are the most highly correlated combinations orthogonal to the first, and so-on.

In the present study, X_1_ and X_2_ represent speech features and the EEG signal respectively. This basic formulation of CCA can be used directly to study the instanteneous interaction between stimulus features and brain response. However, a stimulus at time *t* affects the brain signals for a certain length of time (a few hundreds of milliseconds). Although CCA is a linear approach, simple manipulations of the data allow for its extension to the study of non-linear and convolutional relations, therefore capturing the stimulus-to-brain interaction for a given set of latencies (or time-lags). Here, this is achieved by using a set of filters that capture increasingly long temporal structures (de Cheveigné et al., 2018b). Specifically, we used a dyadic bank of FIR bandpass filters with characteristics (center frequency, bandwidth, duration of impulse response) approximately uniformly distributed on a logarithmic scale. There was a total of 15 channels (N_CH_) with impulse response durations ranging from 2 to 128 samples (2 s). The filterbank was applied to both stimulus and EEG matrices, largely increasing the dimensionality of the data.

Dimensionality reduction was applied to both stimulus and EEG matrices (of size T × N_F_ and T × N_EL_, where T, N_F_, and N_EL_ indicate numbers of time-samples, stimulus features, and EEG electrodes respectively). First, we used PCA and retained N_PC_ < N_EL_ principal components to spatially whiten the EEG data, whose neighbouring channels are largely correlated. The value of this parameter was adjusted using a grid search procedure. Second, the filterbank was applied to both stimulus and EEG data. Finally, PCA was used to reduce the dimensionality of both stimulus and EEG matrices, by retaining N_stim_ < N_F_*N_CH_ and N_EEG_ < N_PC_*N_CH_ components respectively. The CCA models were all trained and tested using a leave-one-out nested cross-validation to control for overfitting. For each outer cross-validation loop, one fold was held-out for testing while a second cross-validation loop was run on the remaining data. In this inner loop, the model hyperparameters were tuned on a held-out validation fold to maximise the sum of the correlation coefficients for the CC-pairs. This framework allowed for the tuning of the values N_stim_ and N_EEG_. In addition, the validation folds at each cross-validation step were used to determine the optimal *shift* between stimulus and neural signals.

#### Temporal Response Function Analysis

Complementary to CCA, a system identification technique was used to compute a channel-specific mapping between each speech representation and the recorded EEG data. This method, commonly referred to as the temporal response function (TRF) analysis (forward model) (Lalor et al., 2006; Ding and Simon, 2012), estimates a filter that optimally describes how the brain transforms the speech features of interest S(*t*) into the corresponding continuous neural responses R(*t*), over a series of pre-specified time-lags: R(*t*) = TRF * S(*t*), where ‘*’ indicated the convolution operator. The TRF values, or weights, were estimated using a regularised linear regression approach, wherein a regularisation parameter was tuned to control for overfitting (Crosse et al., 2016a). One way to use this approach is to study the model weights to identify scalp areas and time-lags that are of particular importance for the specific speech-EEG mapping. A second approach consists of predicting the EEG signals at each channel of interest.

This approach is complementary with CCA analysis in that it provides us with detailed insights on the temporal and spatial patterns. This is possible at the cost of additional constraints, specifically on the frequency-bands of interest and on the magnitude of the prediction correlation values which, since they are calculate in the noisy EEG channel-space (rather than the denoised CCA-space), are usually in the order of 0.05. For this reason, it is preferable to conduct the analysis on the most relevant part of the EEG signals, which can be achieved with a more confined temporal filtering (for an example of the effect of EEG filtering on forward TRF models see Di Liberto et al., 2015). In particular, we restricted the analysis to the frequency-band 0.5-9 Hz (we applied separate low- and high-pass fifth-order Butterworth zero-phase filters).

#### Measuring the quality of the speech-EEG mapping

We used two metrics to quantify the quality of the CCA-based speech-EEG mapping model: correlation and discriminability in a match-vs-mismatch classification task. A Pearson’s correlation coefficient was calculated for each CC-pair. The first CC-pair is the most relevant, but meaningful speech-EEG correlations can arise for an arbitrary number of components. To obtain a measure sensitive to these multiple dimensions, we introduced a match-vs-mismatch classification task that consisted in deciding whether a segment of EEG (duration T_DECODER_) was produced by the segment of speech that gave rise to it, or by some other segment. Discriminability in this task, measured by *d-prime*, reflects the ability of the model to capture the relation between speech and EEG. The *d-prime* metric was derived from the discriminant function of a support vector machine (SVM) classifier trained on the normalised Euclidean distance between pairs of CCs. A cross-validation procedure (*k* = 30) was used in which the classifier was trained and evaluated on distinct data to discriminate between match and mismatch segments. T_DECODER_ was set to the value 1 second, which avoided saturation (classification either too easy or too difficult) in both group and single-subject level analyses.

The quality of the TRF-based speech-EEG mapping was assessed using a correlation metric. Specifically, Pearson’s correlation coefficients were calculated between the EEG signal and its prediction for each scalp electrode separately. This procedure was repeated for TRF models fit using various time-latency windows and stimulus feature-sets, which allowed the pinpointing of latencies that were most relevant to the interaction between EEG and particular speech features of interest (the time-lag windows were within the interval 0 – 900 ms, non-overlapping, and of duration 100 ms). A similar analysis was conducted to investigate topographical patterns corresponding to the various speech-EEG latencies. Specifically, TRF models were fit using a single time-latency window between 0 and 900 ms. Topographical patterns of the corresponding TRF weights were averaged for intervals of interest.

#### Statistical Analyses

Unless otherwise stated, all statistical analyses were performed using two-tailed permutation tests. For tests involving several contiguous time latencies, a cluster-mass non-parametric analysis was conducted, with one as the minimum cluster size (Maris and Oostenveld, 2007). This statistical test takes into consideration the scalp distribution of the measure of interest by performing a permutation test on the cluster of electrodes with the highest score, i.e., the most important cluster according to the metric of interest. This approach provides a solution to the multiple comparison problem by including biophysically-motivated constraints that increase the sensitivity of this statistical test in comparison with a standard Bonferroni correction.

## Results

Non-invasive EEG signals were recorded from ten participants as they listened to an audiobook. We conducted three analyses tackling the questions: 1) Do cortical signals track the small changes in phonotactic probability that characterise natural speech? 2) Can we measure these phonotactic responses at the individual-subject level? And 3) do these signals reflect a pre-lexical influence of phonotactics in speech comprehension?

### Neural Evidence for the Processing of Probabilistic Phonotactics

Brain signals that are common among participants listening to the same speech stimulus were estimated using MCCA (de Cheveigné et al., 2018a). This *consensus signal* (CS) can be thought of as a ground truth cortical response with better signal-to-noise ratio than EEG data of individual subjects. A speech-EEG model based on CCA was then employed to related this consensus EEG signal to different speech feature sets. The quality of the model (measured by correlation and *d-prime* metrics) was used as a measure of the ability of each feature set to capture speech structure predictive of the EEG response.

We wish specifically to evaluate the predictive power of the phonotactic feature set P relative to, and in combination with, other known feature sets such as spectrogram of phonetic features.

We first estimated the quality of a CCA-based model involving only the phonotactic feature vector (P; Figure 1A, **top**) and EEG. The r-value of 0.42 obtained for the first CC-pair was larger than the 99th percentile of a distribution obtained by shuffling the values of the pulses within the P vector while leaving their times intact (median over 100 shuffles: *r* = 0.34; 99^th^ percentile: *r* = 0.35). This result indicates that phonotactic probabilities were reflected by the EEG signals. However the phonotactic feature vector is correlated with other predictive features (such as spectrogram or phonemes), so we cannot be sure that its predictive power stems from phonotactic information per se. For that, we must compare combinations of features that include, or not, the phonotactic vector P. We formed combinations of features including the acoustic spectrogram S (Di Liberto et al., 2015; Lalor et al., 2009; Obleser et al., 2012), a phoneme representation based on phonetic features F (Mesgarani et al., 2014; Di Liberto et al., 2015, 2018a), phoneme onsets O (Brodbeck et al., 2018) and our newly introduced phonotactic features P (see Figure 1A). If each of these features carries information complementary to the others, and not captured by them, we expect speech-EEG correlations to monotonically increase with the inclusion of additional features in the analysis: namely S, FS, OFS, and POFS as schematized in Figure 1B. Indeed, correlation coefficient values for CCA models based on these four combination of features agree with this prediction (Figure 2A; *r_S_* < *r_FS_* < *r_OFS_* < *r_POFS_*). Of possible concern is that these models differ in the number of dimensions (and thus parameters) involved. A large number of parameters can lead to overfitting, which should penalise the models with more features, contrary to what we observe. To further exclude such a possibility, we randomly shuffled the values of the pulses within the phonotactic feature vectors while keeping their timing constant. The distribution of correlation scores for P_shu_OFS obtained by repeated shuffling is indicated in Figure 2A. The value obtained for POFS is above the 99^th^ percentile of that distribution. This same control procedure was applied to the F and O features and confirmed that their respective enhancements are driven by the addition of meaningful features, and not by differences in dimensionality, as they produced stronger correlations than the 99th percentile of the corresponding shuffled distributions. In summary, each of these features carries useful information not carried by the others.

**Figure 1.**
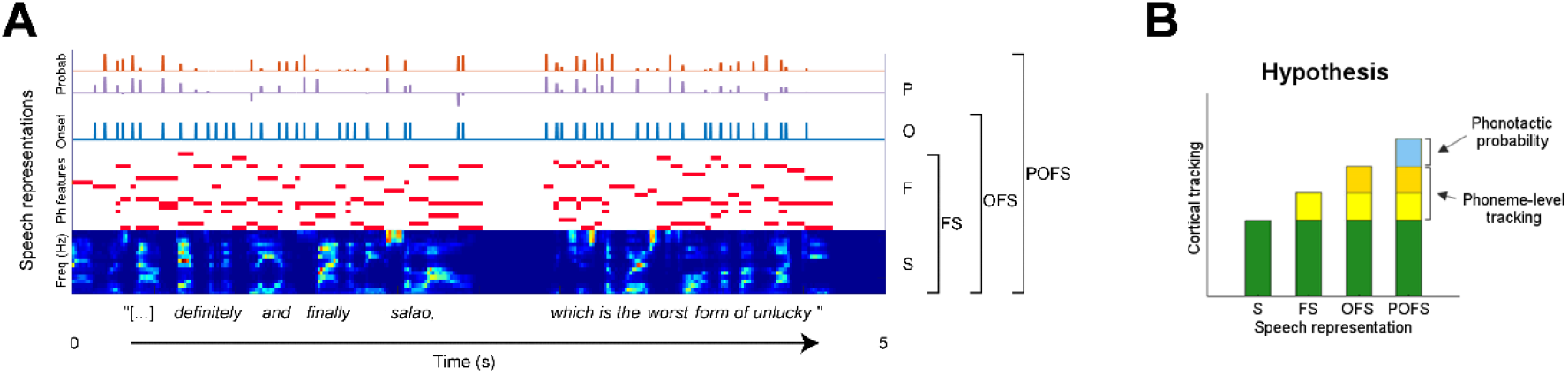
**(A) Speech representations** for a 5 seconds portion of the stimulus. From bottom to top, the acoustic spectrogram (S) which consists of a 16-channel time series of power within 16 frequency bands; phonetic features (F), whose permissible combinations map to English phonemes; phoneme onsets (O), which mark the beginning of each phoneme; and the probabilistic phonotactic vector (P), a representation indicating the inverse likelihood of a sequence (from the beginning of a word to each of its phonemes). **(B) Expected outcomes:** We hypothesise that, if a stimulus representation encodes features not captured by other representations, adding it to the others will improve the prediction of cortical responses. In particular we predict an increase in cortical tracking due when phonotactic probabilities are added to the mix (POFS – OFS, blue increment).

**Figure 2:**
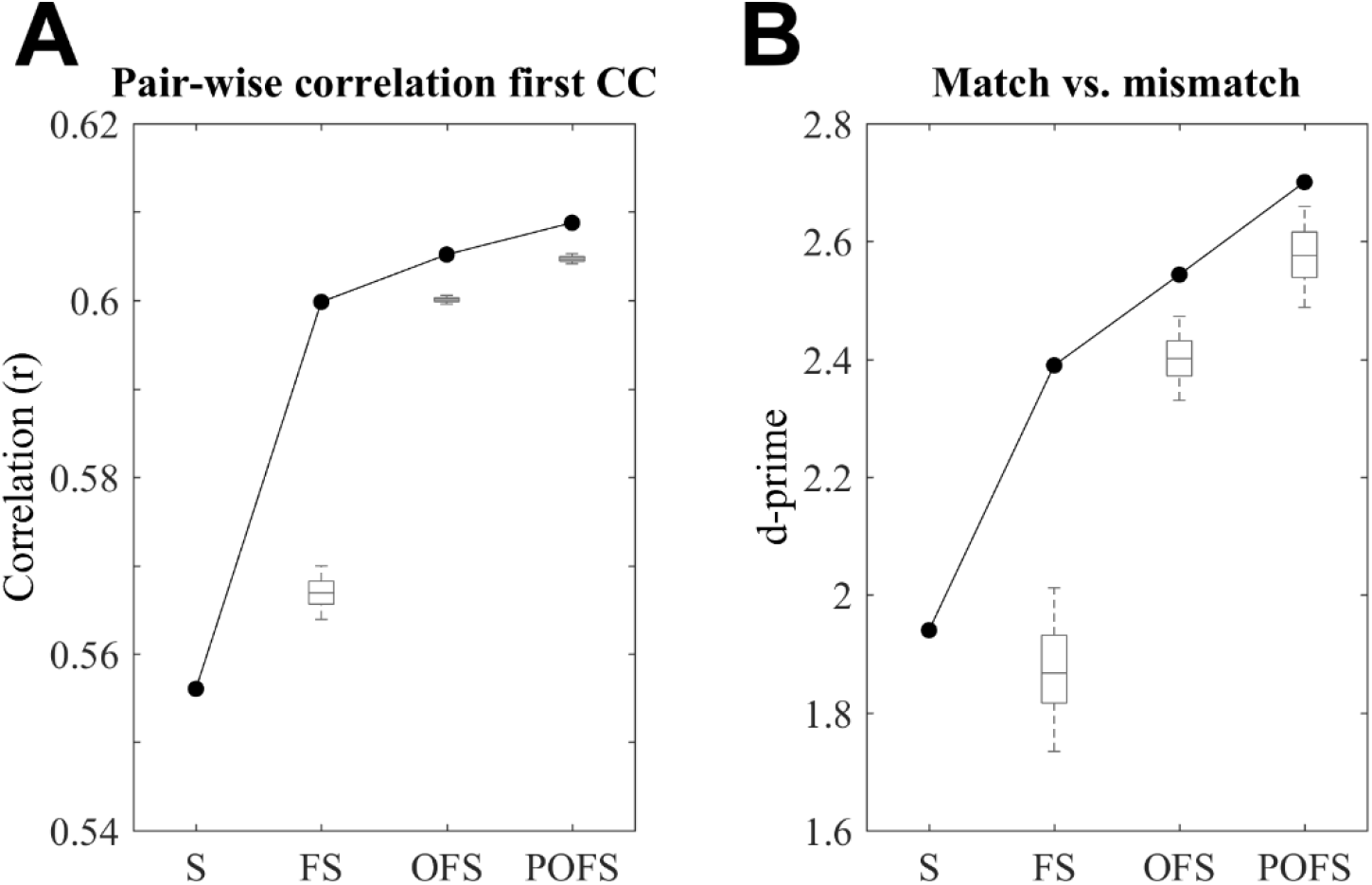
EEG responses to natural speech are best explained when including phonotactic probability among the speech features. Data from all participants were combined using MCCA. This consensus EEG signal (CS) preserves signals that are maximally correlated across subjects. **(A)** A CCA analysis was conducted between each speech representation and the CS signals. Speech-EEG correlations for the first canonical component (CC) pair were best when using the combined model POFS, indicating that phonotactic probabilities explain EEG variance that was not captured by the purely acoustic-phonemic models (S, FS, and OFS). **(B)** In addition, phonotactic probabilities enhanced the *d-prime* score of a match-vs-mismatch classification test. The box-plots indicate the 99^th^ percentile of the performance when using a combined model (FS, OFS, or POFS) after randomly shuffling information for the newly added feature (F, O, and P respectively).

The previous analysis was based on correlations for the first CC-pair only, but other components may carry relevant information as well. To get a more complete picture we performed a similar analysis based on the *d-prime* measure for a match-vs-mismatch trial classification, which combines all components simultaneously (see Methods). The *d-prime* values showed patterns resembling what previously seen for the correlation analysis. Specifically, a *d-prime* of 0.704 resulted from the CCA analysis on P, which was greater than the 99th percentile of the shuffled distribution (median over 100 shuffles: *d-prime* = 0.504; 99^th^ percentile: *d-prime* = 0.544). Furthermore, *d-prime* values monotonically increased for S, FS, OFS, and POFS, showing again greater values than the corresponding shuffle distributions (Figure 2B). The greater value for POFS relative to OFS and P_shu_OFS reinforces our claim that cortical signals track phonotactic probabilities.

### Robust Individual-subject EEG Tracking of Phonotactic Probabilities

The previous analysis provided evidence that the cortical responses to natural speech, measured with non-invasive EEG, are influenced by phonotactic probabilities. To test whether such responses can be reliably measured at the individual-subject level, we conducted the same CCA analysis as in the previous section on the brain recordings from each individual. Figure 3 (left panels of A and B) illustrates both correlation and *d-prime* results. The scores are overall smaller than for the analysis based on the consensus signal, reflecting the greater amount of noise in the subject-specific data, but the same trends are observed. POFS is the best performing model in terms of both correlation (POFS > OFS, *p* = 0.0008; *d* = 1.35; POFS > FS, *p* < 0.0001; *d* = 1.89; POFS > S, *p* < 0.0001; *d* = 1.20) and *d-prime* (POFS > OFS, *p* = 0.027; *d* = 0.60; POFS > FS, *p* = 0.0012; *d* = 0.86; POFS > S, *p* = 0.0049; *d* = 0.98). In addition, this analysis confirmed that phonetic features explain EEG variance not captured by the acoustic spectrogram (FS > S; correlations: *p* = 0.0045, *d* = 1.17; *d-prime*: *p* < 0.014, *d* = 0.85) and, similarly, that the phoneme onsets vector refines the FS representation of speech (OFS > FS; correlations: correlations: *p* < 0.0001, *d* = 1.03; *d-prime*: *p* = 0.042, *d* = 0.65). The average benefits (relative gain) of adding the onset vector O, and the phonotactic vector P, for both measures is plotted in the right-hand panels of Figure 3A and B. Statistical analysis on these average measures confirms that phonotactic information has a measurable effect on the EEG responses to speech (correlation: *p* = 0.0008, *d* = 1.03; *d-prime*: *p* = 0.016, *d* = 0.65).

**Figure 3:**
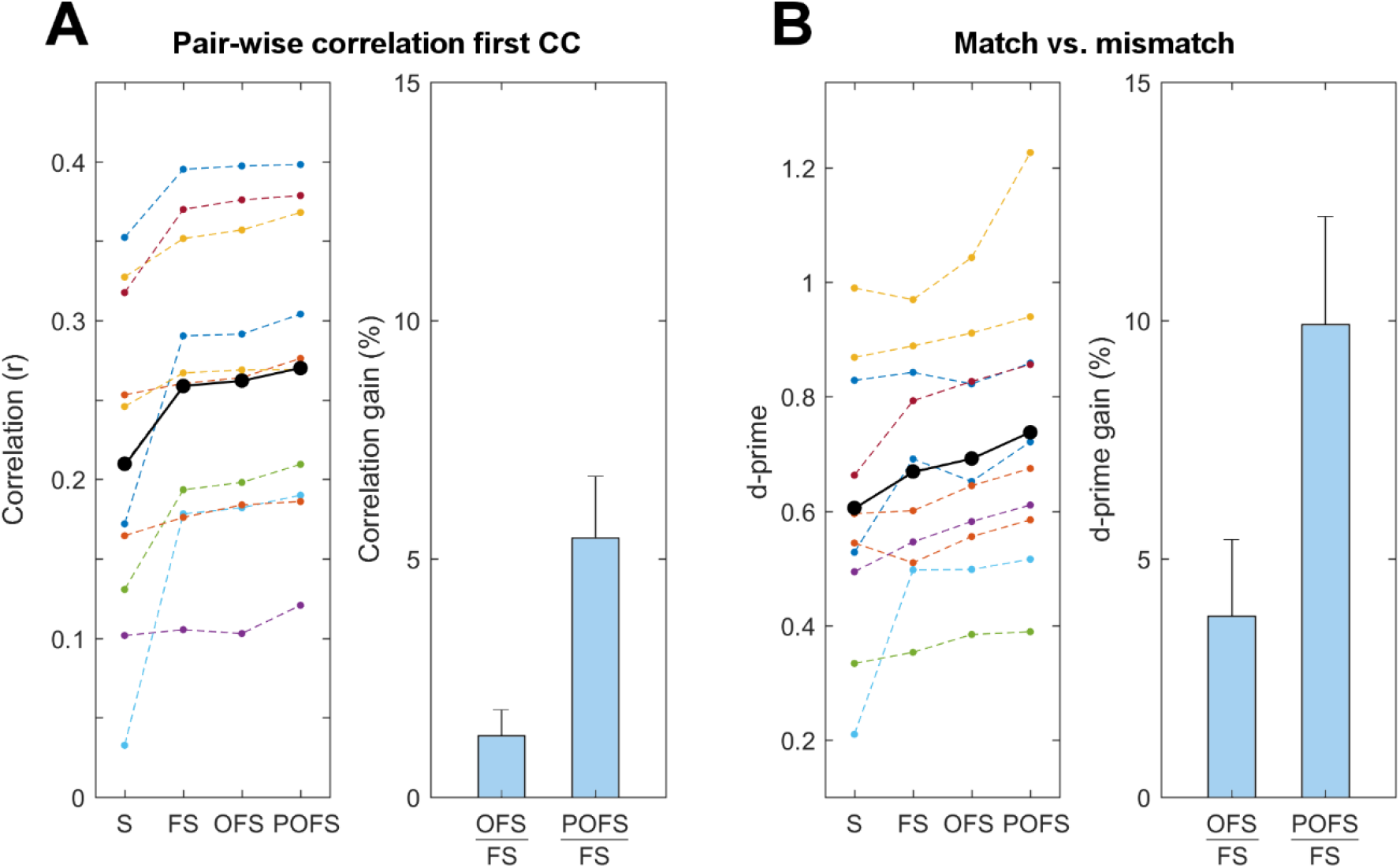
Phonotactic probabilities enhance the speech-EEG mapping at the individual subject level. CCA analyses were conducted between each speech representation and the corresponding EEG responses for each individual subject. **(A)** Speech-EEG correlations for the first canonical component pair were greatest when using the combined model POFS (left panel). The thick black line indicates the average across subjects while the coloured dots/lines refer to the individual subjects. The bar-plot shows the relative correlation gain (%) of the combined models OFS and POFS with FS (i.e. the contribution given by O and P respectively). **(B)** Similar results are shown for the *d-prime* scores of a match-vs-mismatch classification test. Results for individual subjects are colour-coded (same colors as for A). Phonotactic probabilities enhance the single-subject scores for FS and also show significant improvement compared to OFS.

Finally, we conducted additional analyses to test whether other models of phonotactic information can explain EEG responses as well, or better, than P. A first single-subject CCA-based analysis compared P to *neighbourhood density* (P_neigh_). This feature was suggested as a possible neural strategy for an indirect encoding of phonotactic information (Vitevitch et al., 1999; Bailey and Hahn, 2001a). P performed better than this new measure in terms of *d-prime* (POFS > P_neigh_OFS; one-tailed permutation test: *p* = 0.0179; *d* = 0.68). We performed a similar comparison between P and probabilistic definitions of phoneme surprisal (P_sur_) and entropy (P_ent_) (Brodbeck et al., 2018; Gaston and Marantz, 2018). Again, P performed better than these two measures. Specifically, P showed larger *d-prime* values than P_ent_ (POFS > P_ent_OFS; one-tailed permutation test *: p* = 0.037; *d* = 0.67) and P_sur_ (POFS > P_sur_OFS; one-tailed permutation test: *p* = 0.02; *d* = 0.73).

### Timescale of Cortical Responses to Phonotactics

Our results suggests that phonotactic probabilities influence the cortical processing of natural speech. We conducted further analyses to assess the temporal dynamics of this effect. Linear forward models were fit using the TRF approach to describe how speech features are transformed into EEG signals. Because of the sensitivity of the forward TRF method to EEG noise, we restricted the analysis to the frequencies 0.5-9 Hz, which are most relevant for the EEG tracking of speech acoustic and phoneme-level features (Di Liberto et al., 2015, 2018b; Kösem and van Wassenhove, 2016; Vanthornhout et al., 2018).

Forward encoding models were fit for each speech representation (S, FS, OFS, POFS) using non-overlapping time-lag windows of duration 100 ms within the interval 0 – 900 ms. Average EEG prediction correlations confirm the hypothesised general trend that emerged also from the CCA analysis (S < FS < OFS < POFS; Figure 4–1). Crucially, the direct comparison of POFS and OFS reveals a significant effect of phonotactics for a cluster of speech-EEG latencies between 100 and 400 ms (cluster statistics, *p* < 0.05), with peak effect-size at the latency-window 300 – 400 ms (*d* = 1.53) (Figure 4).

**Figure 4:**
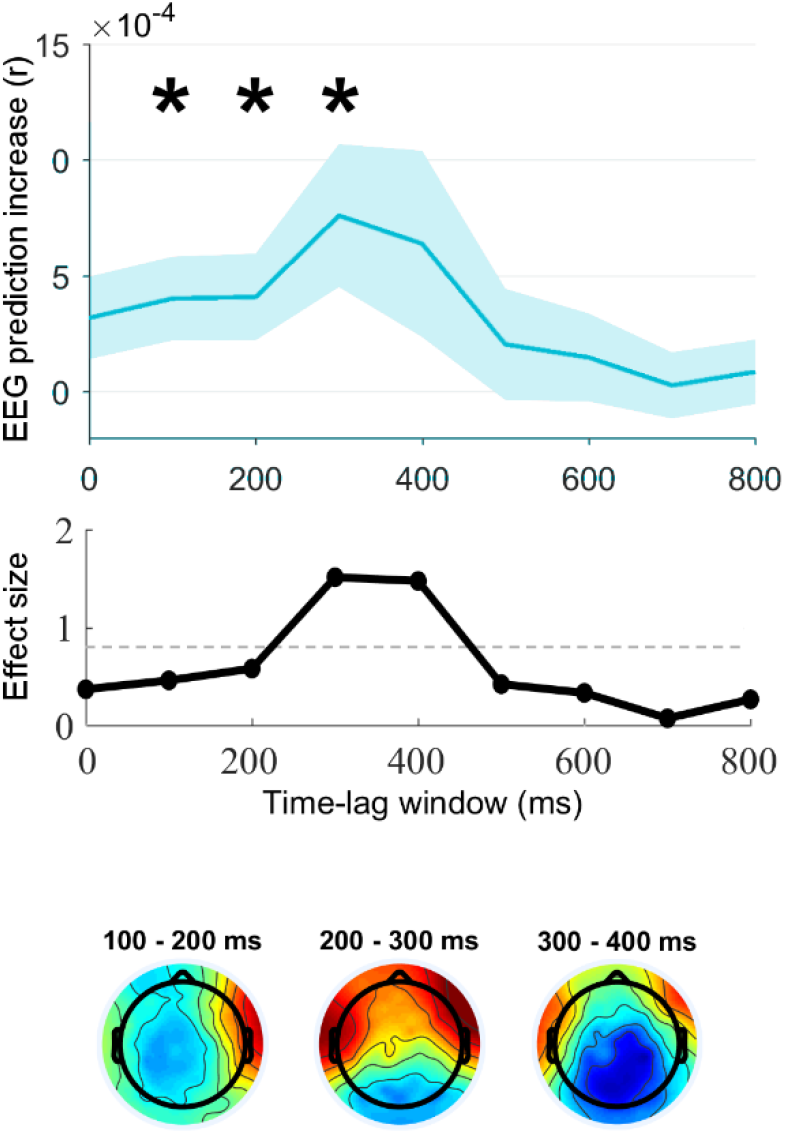
EEG tracking of phonotactic probabilities is specific to speech-brain latencies of 100-400 ms. A temporal response function (TRF) analysis was conducted to estimate the amount of EEG variance explained by phonotactic probabilities for speech-EEG latency windows between 0 and 900 ms and window-size 100 ms. EEG prediction correlations were calculated for different speech feature-sets and for the various speech-EEG latencies. The enhancement in EEG predictions due to phonotactic probabilities is shown for all time-latency windows. Shaded areas indicate the standard error of the mean (SE) across subjects. Stars indicate significant enhancement (* *p* < 0.05) as a result of a cluster mass statistics (top). Cohen’s *d* was calculated to measure the effect size of the enhancement due to phonotactics. Values above 0.8 are considered as ‘large’ effects (above dashed grey line) (centre). Topographical patterns of the TRF weights for a model fit over time-lags from 0 to 900 ms are shown for latencies with a significant effect of phonotactic probabilities (100-400 ms) (bottom).

## Discussion

Our results demonstrate that cortical responses to natural speech reflect probabilistic phonotactics. First, linear modelling revealed a time-locked interaction between phonotactic information and low-frequency EEG. Then, we established that brain responses to phonotactics can be measured at the individual subject-level. Finally, we found that speech-EEG latencies of 100-400 ms are most relevant to those brain responses, suggesting that phonotactic information contributes to natural speech processing at pre-lexical stages.

### A novel measure of phonotactic processing

Phonotactic information plays an important role in speech perception. However, crucial questions remain unanswered about the underpinnings of the corresponding cortical processes, mainly due to a lack of tools to extract direct measures of brain responses to phonotactics. Although neurophysiology has partially fulfilled this need (Connolly and Phillips, 1994; Dehaene-Lambertz et al., 2000; Wagner et al., 2012; Cibelli et al., 2015; Leonard et al., 2015), its findings were mainly confined to nonsense words or to the domain of phonotactic violations, which are exceptions in natural speech scenarios. The present study aimed to measure brain signals corresponding to the continuous integration of phonotactic information, which are difficult to isolate when measuring only phonotactic violations. These violations trigger various other processes such as phonological repair, which may emerge in the evoked-response (Dehaene-Lambertz et al., 2000; Dupoux and Pallier, 2001; Domahs et al., 2009). Here, we found evidence that cortical responses to narrative speech reflect the well-formedness of phoneme segments as expressed by probability values, therefore with no (or very few) violations and phonological repair (Figures 2 and 3). This finding pushes beyond the phonotactic violation paradigm and provides us with a tool based on linear models to isolate measures of phonotactic-level processing during natural speech perception.

This work constitutes a further step towards the characterisation of brain responses to natural speech, adding to recent work aimed at isolating brain responses to distinct processing stages, involving speech acoustics (Ding and Simon, 2014), phonemes (Di Liberto et al., 2015, 2018c), sentence structure (Ding et al., 2015, 2017), and semantic similarity (Broderick et al., 2018). The ability to simultaneously account for and disentangle brain responses to continuous speech at different processing stages constitutes a novel and powerful tool to study the neurophysiology of speech. In particular, isolating brain responses to phonotactics could provide new insights on the positive impact of this mechanism in case of language impairment, and also when the phenomenon plays against us. For example, when learning a second language, these brain mechanisms cause misperception and mispronunciation, and contribute to stereotypical accents (Davidson, 2006a, 2006b; Lentz and Kager, 2015). In addition, the present framework produces objective measures indicating how strongly EEG responses to speech correspond with a particular phonotactic model, thus offering a new opportunity to test the neurophysiological validity of theoretical and computational models (e.g. BLICK).

Our results provide new insights in this direction, indicating that phonotactic probabilities, as defined by the computational model BLICK, are better represented in the EEG signal than a purely probabilistic definition of phoneme probability (P_sur_, P_ent_) (Gaston and Marantz, 2018) and, importantly, than phonological neighbourhood density (P_neigh_) (Vitevitch et al., 1999; Frisch et al., 2000; Bailey and Hahn, 2001a). While further studies could explore other hypotheses on the encoding and processing of phonotactic information more comprehensively, the present finding is in line with research suggesting distinct roles for phonotactics and neighbourhood density (Vitevitch et al., 1999; Bailey and Hahn, 2001a; Storkel et al., 2006). Specifically, the first would aid speech perception by facilitating processing and triggering learning of new words at early pre-lexical stages, while the latter would influence the integration of new and existing lexical representations at a later stage.

A similar issue relates to the speech-brain latencies associated with phonotactic information. Indeed, previous research found interactions between phonotactic violations and evoked brain components such as N400 and LPC (Domahs et al., 2009; White and Chiu, 2017). It has also been suggested that the N400 magnitude may be directly linked to phonotactic information, while effects at the longer LPC latencies may be spurious and, instead, reflect other related processes, such as changes in cognitive load related to the size of the neighbourhood of permissible words (Dupoux et al., 1999; Vitevitch et al., 1999; Dupoux and Pallier, 2001; Storkel et al., 2006). Our results contribute to this debate by suggesting that latencies of 100-400 ms are the most relevant for the processing of phonotactic probabilities. Furthermore, topographical patterns at those latencies present activations over centro-parietal scalp areas that qualitatively resemble that of an N400 component. One possibility is that this response is related to an early N400, whose latencies reflect the rapid processing of phonotactics in a natural speech scenario. It is also possible that this response reflects multiple cortical correlates, one in correspondence with the earlier weaker effect (100-300 ms) (Brodbeck et al., 2018), and a separate one with a larger effect-size at longer latencies (300-400 ms) (Pylkkänen et al., 2002, 2000). A direct comparison between EEG responses to phonotactic probabilities and phonotactic violations could clarify some of these issues, as previously attempted in the similar context of semantic-level processing (Broderick et al., 2018).

### Theoretical implications of a rapid time-locked response to phonotactics

Our results have important implications for current theories on phonotactics, by providing insights into both temporal dynamics (when) and neural encoding (how) of this cortical mechanism. Phonotactic information, which aids speech recognition and learning of new words (Mattys and Jusczyk, 2001; Munz, 2017), was suggested to involve one of the following: 1) the phoneme identification stage (one-step models; Dehaene-Lambertz et al., 2000; Dupoux et al., 2011); 2) a pre-lexical stage that occurs after phoneme identification (two-step models; Church, 1987); or 3) a later lexical stage that influences pre-lexical processes through feedback connections (lexicalist models; McClelland et al., 2006; McClelland and Elman, 1986). In this context, a large body of literature in psycholinguistics supports a pre-lexical account of phonotactics (McQueen, 1998; Jusczyk et al., 1999; Sebastián-Gallés, 2007). For example, infants showed sensitivity to phonotactics by 9 months of age, suggesting that this information aids speech segmentation even at early developmental stages, before being able to understand speech (Jusczyk et al., 1994). Similarly, it was shown that humans are sensitive to phonotactic information even when meaning is not involved (nonsense words), pointing to the early implementation of phonotactic repair (Dupoux et al., 1999; Davidson, 2011; Rossi et al., 2013). This indirect evidence for a pre-lexical influence of phonotactic information finds experimental support in both phonotactic violation studies (Dehaene-Lambertz et al., 2000; Pylkkänen et al., 2002) and in the present work, which isolated cortical responses to probabilistic phonotactics showing short speech-EEG latencies (100-400 ms).

Indeed, it is possible that other post-lexical brain responses to phonotactics exist but could not be measured. In fact, such higher-level effects could exhibit weaker time-locking, which would hamper the ability to capture them with our framework. Indeed, this hypothesis should be tested with more controlled experimental paradigms, possibly by making less assumptions on the time-locking between phonotactics and brain signals. Although we cannot be conclusive on this point, the latencies of 100-400 ms could be in line with one-step models (Dehaene-Lambertz et al., 2000; Dupoux et al., 2011), which hypothesise that phonotactic processing occurs pre-lexically and together with phoneme identification, whose EEG responses were measured for latencies up to 300 ms (Di Liberto et al., 2015; Khalighinejad et al., 2017).

In summary, our results indicate rapid time-locked brain responses to probabilistic phonotactics. This phenomenon emerged for low-frequency cortical signals (< 9 Hz) and were reliably measured at the individual subject-level. We also found that the speech-EEG latencies of 100-400 ms most strongly reflects phonotactic information, which is in line with a pre-lexical account of phonotactic processing. This provides the field with a new tool to study the brain processing of phonotactics using natural speech.

## Author Contributions

G.D.L. conceived the study and collected the data. G.D.L., A.d.C., and D.W. formulated the data analysis procedure. G.D.L. analysed the data. G.D.L. wrote the first draft of the manuscript. A.d.C., G.A.M., and D.W. edited the manuscript.

## Acknowledgements

This study was supported by the EU H2020-ICT grant 644732 (COCOHA). The authors would like to thank Dorothée Arzounian for useful discussions at the start of this study.

## Extended data

**Figure 4-1.**
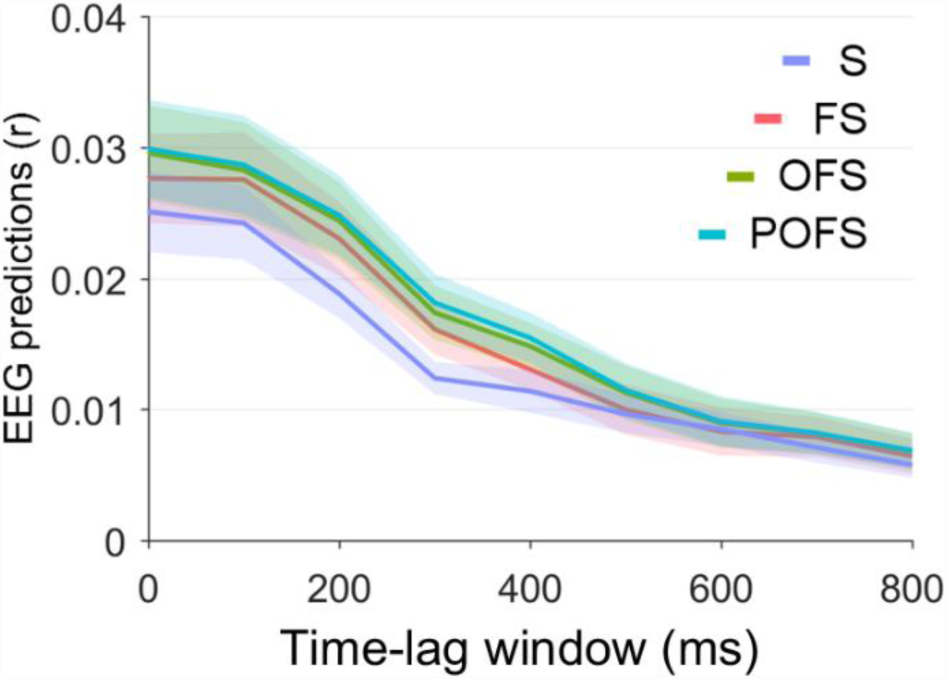
A temporal response function (TRF) analysis was conducted to estimate the amount of EEG variance explained by phonotactic probabilities for speech-EEG latency windows between 0 and 900 ms and window-size 100 ms. EEG prediction correlations averaged across all scalp electrodes are shown for different speech feature-sets and for the various speech-EEG latencies. Shaded areas indicate the standard error of the mean (SE) across subjects. The contrast between EEG prediction values for POFS and OFS is shown in Figure 4.

## Notes

Conflicts of interest: none declared.

